# Rac negative feedback links local PIP_3_ rate-of-change to dynamic control of neutrophil guidance

**DOI:** 10.1101/2022.12.30.521706

**Authors:** Jason Town, Orion Weiner

## Abstract

To migrate efficiently, neutrophils must polarize their cytoskeletal regulators along a single axis of motion. This polarization process is thought to be mediated through local positive feedback that amplifies leading edge signals and global negative feedback that enables sites of positive feedback to compete for dominance. Though this two-component model efficiently establishes cell polarity, it has potential limitations, including a tendency to “lock” onto a particular direction, limiting the ability of cells to reorient. We use spatially-defined optogenetic control of a leading edge organizer (PI3K) to probe how cells balance “decisiveness” needed to polarize in a single direction with the flexibility needed to respond to new cues. Underlying this balancing act is a local Rac inhibitor that destabilizes the leading edge to promote exploration. We show that this local inhibitor enables cells to process input signal dynamics, linking front stability and orientation to local temporal increases in input signals.

## Introduction

Neutrophils find and kill invading pathogens by dynamically aligning their front-back polarity axis with subtle external gradients of chemoattractants that indicate injury or infection. One of the fundamental problems these cells need to solve is the consolidation of the biochemical signals that drive protrusions to a small portion of the cell surface. To navigate their complex environments efficiently, migrating cells must balance this “decisive” consolidation process with the “flexibility” needed to continuously update the direction of this polarity axis.

Cells consolidate protrusive activity through a reaction-diffusion Turing system (1) that combines local positive and global negative feedback (2–6). In neutrophils, the positive feedback loop is linked to protrusion signaling and involves phospholipids such as PIP_3_, small GTPases such as Rac, and f-actin (7–9). These protrusions push on the membrane, generating increases in tension that rapidly equilibrate throughout the cell (10) and stimulate global inhibition of Rac activation and actin polymerization (5, 11). This model explains several key cell behaviors, including the consolidation of protrusive activity into a single front and the ability to detect chemoattractant gradients over a wide range of ambient concentrations. However, this model has a fundamental limitation: Although simulated cells can properly orient to an initial gradient, they tend to “lock” and have difficulty reorienting their polarity when correcting errors in orientation or when the external gradient changes (3).

To resolve this limitation, Meinhardt proposed the existence of an additional factor, a local inhibitor, that slowly destabilizes fronts of migrating cells, preventing “front-locking” (3). Modeling approaches have made use of local inhibition to produce realistic simulated chemotactic behaviors, such as pseudopod-splitting in *Dictyostelium* (12). Modeling can be used to show how these signaling topologies could function even without knowledge of the specific molecular components. With suitable inputs, the action of a local inhibitor can be demonstrated experimentally even if the specific molecular candidate is not known (13). Experimental studies have advanced our understanding of several potential molecular candidates of a Meinhardt-style local inhibitor (14–18). However, it is unclear whether these candidates act locally due to possible confounding effects from other globally acting inhibitors. To formally demonstrate the existence of a local inhibitor, these confounding effects would need to be removed from the system (19), but this typically disrupts the organization of the front signals (20–22).

Here we overcome this challenge by using an optogenetic approach to control the production of PIP_3_, a key protrusion-activating signal in migrating neutrophil-like HL-60 cells (7, 8, 23, 24). Using this approach in combination with dynamic, computer-controlled spatial stimulation and pharmacological perturbations, we demonstrate the existence of a local negative feedback loop operating on Rac at the leading edge of migrating neutrophils. We show that this local negative feedback loop allows cells to detect the local rate of signal increase, consistent with the pilot pseudopod model for cell guidance first described by Gerisch (25). We modify the basic pilot pseudopod model, accounting for insensitivity at the backs of cells (21, 23, 26–29) and for the basally-higher levels of PIP_3_ at the fronts of migrating cells. These two modifications explain the particular sensitivity of the lateral edges of migrating cells to PIP_3_ signals. The edges are within the sensitive region and have more “room to grow” compared to the center of the front. This “peripheral attention” model describes how cells use temporal features of internal signals to continuously steer their fronts and balance decisiveness and flexibility during navigation.

## Results

### Local opto-PI3K stimulation steers migrating HL-60 cells

Previous work has found that local activation of PI3K signaling induces the generation of actin-based protrusions such as pseudopodia and growth cones (10, 24, 30). Since polarized cells differ in the biochemical compositions of their fronts and backs, we first sought to understand whether the response to local PI3K activation spatially varied in migrating HL-60 cells. Towards this end, we used an optogenetic approach to control PI3K signaling (opto-PI3K) in time and space. This enabled us to probe migrating cells’ biochemical and directional responses to user-defined changes in PIP_3_ at different subcellular locations.

Our opto-PI3K approach uses an iLID system (31) to recruit endogenous PI3K to the plasma membrane of HL-60 cells in response to blue light **(Fig. 2A)**. We verified the effectiveness of our optogenetic system using spatially patterned blue light stimulation (470 nm) and PHAkt-Halo, a live-cell biosensor for PIP_3_ production (32–34) **(Fig. S1; Movie S1)**. Upon local activation of PIP_3_ production, we observe a rapid local increase in the localization of Pak-PBD-mCherry. This livecell biosensor predominantly recognizes active Rac in HL-60 cells (35) and typically localizes to the fronts of migrating cells. Inspired by previous work in smart microscopy control systems (36–40), we implemented a computer-vision-based feedback system that tracks the dynamic features of migrating cells (primarily location, polarity, and visibility in TIRF) and delivers automatically updated, spatially-patterned optogenetic stimuli **(Fig. 2B)**. Using this system, we can reproducibly track and perturb PIP_3_ signals in time and space in moving cells.

Local opto-PI3K stimulation at a given subcellular location (front, side, back) caused reproducible behaviors across migrating cells **(Fig. 2C; Movie S2)**. Local activation at the fronts of cells caused highly directed movement in the direction of stimulation, and this motion was more directed than either unexposed control or globally stimulated cells. Local activation at the lateral edges of cells caused cells to turn toward and persistently migrate in the direction of the optogenetic stimulus. Local activation at the backs of cells caused them to either perform a “u-turn” (41) or depolarize, in which case the cell footprint disappeared from the TIRF plane. The directional stimuli generally caused continuous turning responses as opposed to a depolarization/repolarization response. We could even cause cells to turn continuously by updating the position of the stimulus to continuously target one edge of the cell (**Movie S3**). With this controllable perturbation, we now had the ability to ask which spatial and temporal features of PIP_3_ signaling contributed to these cell behaviors.

### Sensitivity to local PIP_3_ stimulation is spatially biased to the lateral edges in polarized cells

We next sought to determine whether cell responsiveness to optogenetic stimulation and downstream activation varied spatially in polarized cells, which are known to differ in the biochemical compositions of their fronts and backs. As other researchers have observed with various signaling inputs (21, 23, 26–29), we found the backs of cells to be relatively insensitive to stimulation via opto-PI3K. Cells took longer to reorient in the direction of the stimulus and displayed u-turn behaviors in response to stimulation at the back. Restricting signaling competency to the fronts of cells could be one mechanism to ensure persistent movement, though, at an extreme, this strategy could cause polarity locking (3). To quantify the spatial extent of this bias in cells moving in an approximately two-dimensional environment, we took advantage of our ability to simultaneously stimulate multiple locations in moving cells using our computer-vision-driven system **(Fig. 2D)**. This allowed us to expand upon results from previous work on back-insensitivity that constrained cell movement in one-dimensional channels (28, 29).

When we activated both lateral edges of a migrating cell in a 180°-opposed fashion, cells stably turned toward only one of the two stimuli **(Fig. 2G, Left; Movie S4, Left; Fig. S2, Left)**. When the opposed stimuli were oriented perpendicularly relative to the initial direction of motion, cells broke symmetry evenly, with half of the cells migrating to the left and the other half migrating to the right. By varying the angle of these opposed competing signals relative to the initial polarity axis, we found that cells were more likely to turn toward stimuli closer to the existing front **(Fig. 2D)**. At the extreme, all cells stimulated simultaneously at their front and back continued moving in their initial direction. These results support the notion that the back of the cell responds less strongly to our optogenetic input.

Cells stably moved toward only one of the two optogenetic stimuli, indicating that this assay created a winner-take-all scenario. Surprisingly, however, the frontward advantage was gradual. In a straightforward winner-take-all system, such as one made from coupled local positive feedback and global negative feedback, any slight initial advantage should be amplified, leading to switch-like advantages. Thus, we might have expected cells to reliably turn toward more front-oriented signals, even for small angles. We sought to resolve this apparent contradiction by further exploring the winnertake-all nature of two-spot competition assays.

### Fronts of polarized cells show behaviors consistent with local negative regulation

The observed stability of migration decisions in response to 180°-opposed stimuli could represent an intrinsic ability of cells to commit to one site of PIP_3_ stimulation by suppressing signaling elsewhere. Alternatively, it could represent an initial commitment that becomes stable due to the insensitivity of the back of the cell to stimulation. To differentiate between these possibilities, we altered the stimulation assay to ensure that both stimulation sites would be continuously near the fronts of cells. We accomplished this by placing activation spots at the cell edges +/-45 degrees from the polarity axis **(Fig. 2G, Right; Movie S4, Right; Fig. S2, Right)**. In this assay, if cells were to direct their fronts toward one stimulus, the other stimulus would be located at the lateral edge of the cell (which should be sensitive to stimulation). During the 5 minutes of frontward-oriented opposed-stimulus exposure, 48% of cells showed some degree of reversal between the two orientations (compared to 6% in the 180°-opposed stimuli). This result suggests that the insensitivity of the backs of cells drives the previously observed stability in the opposed spot competition assay. The long-term stability is not due to an intrinsic ability to suppress signaling at a distance. On the contrary, the orientation switching suggests that some factor locally inhibits front signaling, destabilizing fronts on a slow timescale and making them less able to suppress signaling at distant sites over time.

One additional piece of evidence in support of a local front inhibitor is the difference in directional responses of the cells to local front stimulation and global stimulation assays **(Fig. 2E)**. In the global stimulation assay, cells tend to deviate from their initial direction significantly more than cells driven to migrate forward via local optogenetic stimulation. Local inhibition at the center of the fronts of cells could explain this behavior by resulting in larger responses at regions just outside this area.

This front-localized negative regulation is also supported by measuring the local biochemical responses to global stimulation **(Fig. 2F)**. By quantifying the increases in PIP_3_ and Rac biosensor recruitment to the TIRF plane in cells in response to global stimulation, we observe that the largest fold-change for both PIP_3_ and Rac biosensor recruitment occurs 3-5 µm away from the point at the center of the leading edge of the cell. The combination of insensitivity at the backs of cells (21, 23, 26–29) and local-inhibition-based insensitivity at the extreme fronts of cells could explain this peak.

Our data suggest the existence of local inhibition at the fronts of cells, as predicted by Meinhardt (3). This front-based inhibition is separate from previously observed inhibition (or lack of responsiveness) at the backs of cells and is instead expected to enable directional plasticity. To directly demonstrate local negative regulation at the fronts of cells, we next sought to remove confounding effects from global negative feedback.

### Direct demonstration of local negative feedback for Rac activation

In HL-60 cells with an intact cytoskeleton, protrusion generation causes rapid actin-dependent increases in membrane tension throughout the cell (10); thus, we expect cells to limit protrusion growth via mechanically-mediated global negative feedback (5, 11, 42, 43). Since both global and local inhibition act on and are acted on by similar cues, they are difficult to disambiguate. To isolate the effects of local inhibition, we blocked actin-based protrusions with latrunculin, preventing activation of mechanically-gated global inhibition. Inhibiting this particular node is also expected to interrupt positive feedback (8) **(Fig. 1B)**, leaving only the putative influence of local negative regulation **(Fig. 3A)**.

**Fig. 1.**
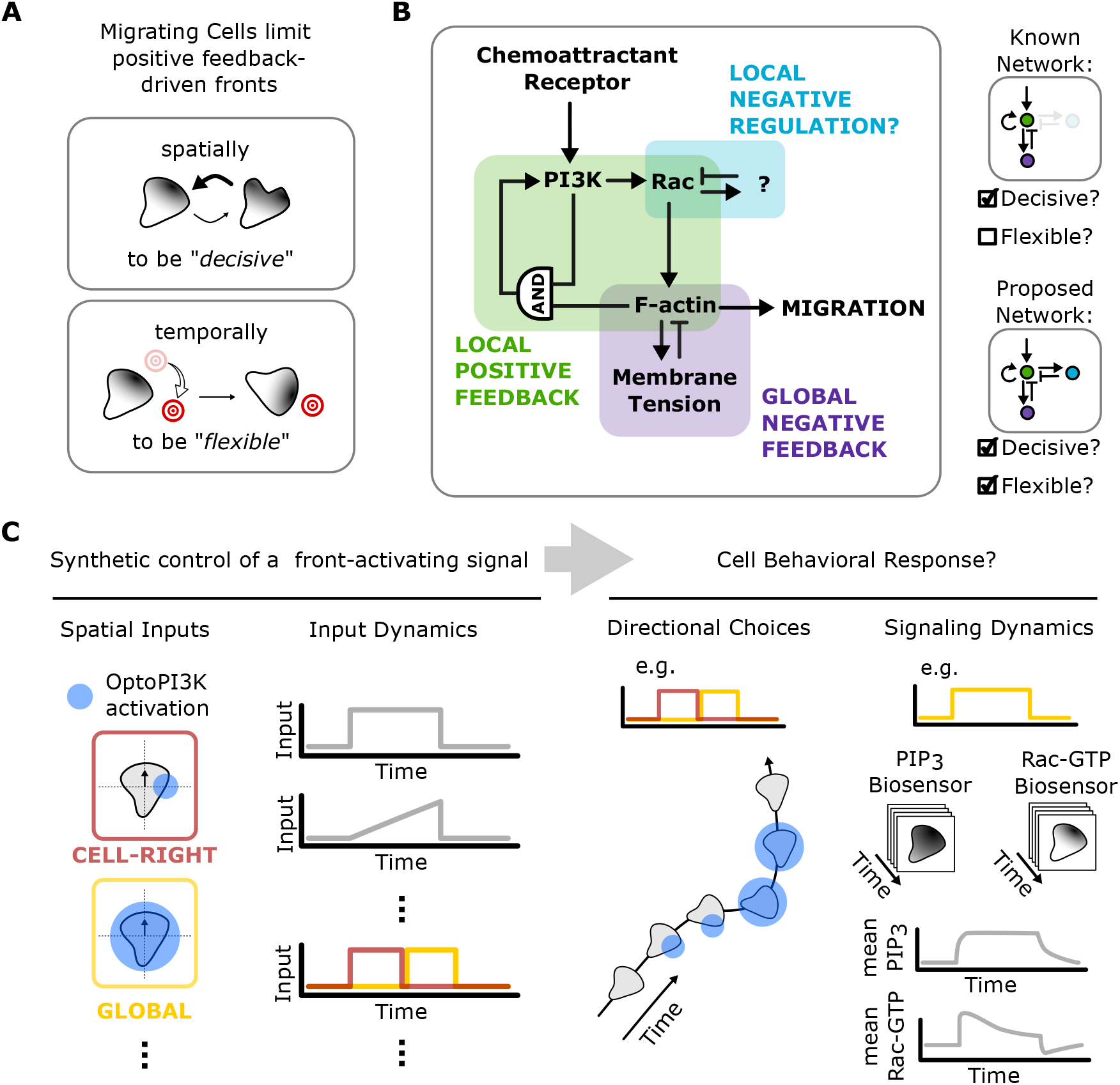
Approach for probing how rapidly migrating cells balance decisiveness and flexibility. **A)** Migrating cells use positive feedback to polarize and build fronts. This positive feedback must be constrained by negative feedback to enable generation of a singular front that can be reoriented. **B)** Previous work has established a central role for local positive feedback and global negative feedback in cell polarity and movement. Models that include only these two feedback loops tend to have “front-locking” behaviors where they become insensitive to new stimuli once polarized. Thus they are decisive in their ability to polarize, but inflexible. One potential solution to the front-locking behavior is the addition of local negative feedback to destabilize the front. Systems including local positive feedback and both local and global negative feedback have the potential to be both decisive and flexible. **C)** Direct spatiotemporal control over a front-activating signal provides a powerful strategy for better understanding how positive and negative feedback modules interact to regulate decision-making in migration.

To avoid previously observed oscillatory dynamics of Rac activity in latrunculin-treated differentiated HL-60 cells (20– 22), we elected to use undifferentiated HL-60 cells, which retain the ability to generate protrusions (10) in response to opto-PI3K stimulation, but are basally quiescent. Latrunculin treatment did not interfere with the ability of our optogenetic tool to generate PIP_3_ and subsequently activate Rac in this context **(Fig. 3B)**. Upon continued, constant local activation of PIP_3_, as read out by our PIP_3_ reporter PH-Akt, we observed a transient activation and then a decline in Rac activity as read out by our Rac biosensor Pak-PBD. We confirmed this transient Rac activation using an ELISA-like Rac activation assay (Rac 1,2,3 GLISA assay, Cytoskeleton Inc) in latrunculin-treated cells **(Fig. S3)**.

The transient Rac response in latrunculin-treated cells suggests a mode of actin-independent inhibition. To determine the spatial range of this inhibition, we subjected cells to a secondary global stimulus following local activation. When we stimulated a small portion of the cell, we observed a transient increase in the localization of the Rac biosensor. When we next allowed the cell to recover briefly in the absence of optogenetic input, a subsequent global stimulation elicited an asymmetric Rac response that was stronger on the previously unstimulated side of the cell compared to the pre-stimulated side **(Fig. 3C; Movie S5)**. Though the timing of the pulses differed slightly between experiments, we consistently observed a spatial reversal in the direction of the Rac response following global stimulation. Furthermore, by applying temporally spaced global pulses, we identified a 90s recovery half-time of the response **(Fig. S4)**. Together these data suggest that local PIP_3_ generation leads to recruitment of a reversible, locally acting, moderately persistent Rac inhibitor.

### Local inhibition operates through negative feedback, linking front signaling to input rates

Beyond its potential role in avoiding the locking of cell polarity decisions, we examined how local Rac inhibition could influence the interpretation of guidance cues. Bacteria chemotax by using integral negative feedback to temporally sense spatial signal gradients while they move through them (46, 47). The internally-measured rate of change of the external signal is used to bias tumbling frequency and accomplish a biased random walk (48). We wondered whether the local front-based inhibition we had identified in neutrophils might play an analogous role in their interpretation of guidance cues by enabling local temporal sensing. To investigate this possibility, we took advantage of the titratability and controllability of our system to understand how PIP_3_ input dynamics regulate the Rac response. If Rac operates through an integral negative feedback circuit, it should respond to the rate of change in PIP_3_ (analogously to (49, 50)); A rapid increase in PIP_3_ should generate a significant transient increase in Rac activation, whereas a slow ramp of PIP_3_ to the same concentration should generate little to no Rac activation, and this is precisely what we observe **(Fig. 4A)**.

Rate-sensitive responses are a general feature of negative-feedback-based adaptive systems, and the Rac activity response to differing PIP_3_ input dynamics is consistent with adaptation through negative feedback rather than through another mechanism like incoherent feedforward (44, 45, 51– 53). To further evaluate whether our system is operating through negative feedback, we tested the effect of mild Rac inhibition on the dynamics of the response. If the system operates through a Rac-dependent negative feedback loop, then mild inhibition of Rac should disrupt that negative feedback and impair adaptation. By partially inhibiting the ability of Rac to signal to its downstream effectors with mild EHT1864 treatment (54), we observed a dose-dependent shift from transient to sustained Rac activation following an opto-PI3K step input **(Fig. 4B)**. These data suggest that Rac activity is required for Rac inhibition, consistent with Rac-dependent negative feedback.

By operating through adaptive negative feedback, a Meinhardt-style local inhibitor could use rates of local input signal changes (as opposed to the absolute magnitude of input signals) to regulate protrusion lifetimes. This rate dependence is closely related to the pilot pseudopod model for gradient interpretation, as initially proposed by Gerisch (25). This model proposes that migrating cells can navigate chemoattractant gradients across many orders of magnitude by maintaining (randomly initiated) pseudopodia that experience a temporal increase in signal as they extend up a gradient at the expense of those that experience a temporal decrease in signal when extending down a gradient. We next sought to investigate whether local changes in PIP_3_ levels regulate cell guidance, even in the absence of PIP_3_ gradients across the front.

### Fronts of polarized cells respond to input signal rates, consistent with a pilot pseudopod model

Our work demonstrates that local negative feedback makes biochemical front signals in cells sensitive to temporal changes in signaling inputs, consistent with a pilot pseudopod model of gradient interpretation. We returned to using our computer-vision-based stimulation assays in moving cells to extend our biochemical signaling observations to cell guidance decisions. We sought to disambiguate the role of temporal changes in input signals from their absolute levels by performing an experiment analogous to our shadow experiment **(Fig. 3)**. Since the backs of cells are relatively insensitive to stimulation, we focused on stimulation of the lateral edges of migrating cells. When we persistently activate cells with local opto-PI3K stimulation at one lateral edge, we can direct them to continuously turn clockwise (for right edge stimulation) or counterclockwise (for left edge stimulation, see **Movie S3**). By switching cells from local to global activation, both edges of the cell attain the same final level of signaling but through different histories. The pre-stimulated edge will have a smaller increase than the stimulus-naive edge **(Fig. 5A)**. If cells respond to signaling levels without history dependence, they should respond to the global stimulus as they did previously **(Fig. 2E, blue traces)**. In contrast, if cells respond to temporal changes in input signals (such as through the local Rac negative feedback loop we have demonstrated), we expect cells to reverse directionality due to the larger increase on their stimulus-naive sides. Our experiments confirm the local temporal interpretation of PIP_3_ inputs **(Fig. 5B; Movie S6)** in a manner analogous to our shadow experiments in latrunculin-treated cells **(Fig. 3C; Movie S5)**.

**Fig. 2.**
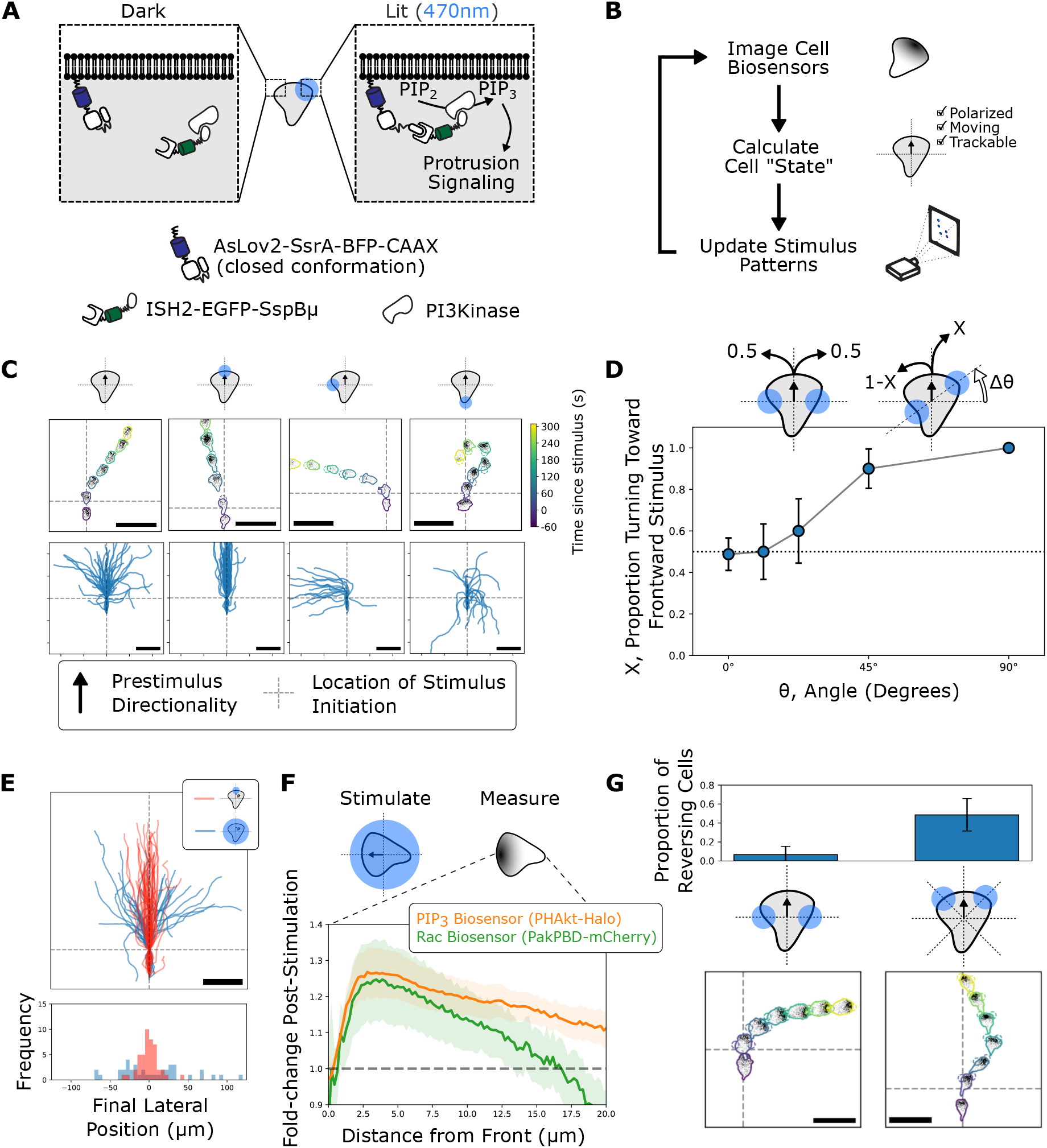
Optogenetic control of PI3K signaling in migrating cells reveals spatial differences in responsiveness to input signals **A)** Schematic of the optogenetic system for controlling PIP_3_ generation. **B)** Computer-vision-based strategy for automatically updating optogenetic input for moving cells. **C)** Local opto-PI3K stimulation at the fronts, sides, or backs of migrating cells causes corresponding changes in cell directionality. See **Movie S2** for an animated version of this data. Scale bars: 50 µm. **D)** When cells are stimulated at two opposing sites of opto-PI3K activation, they break symmetry and decisively move toward one of the two inputs. When the angle of these opposed sites of stimulation is varied relative to the initial polarity of the cell, a preference for the frontward-directed input is revealed. **E)** The frontward-bias shown in D does not cause globally-stimulated cells to persist in their current direction of movement. Scale bars: 50 µm. **F)** Fold-change in biosensor recruitment as a function of font-to-back distance in response to global stimulation shows a peak response for Rac activation 3 to 4 microns away from the leading edge of the cell in response to opto-PI3K input. **G)** Cells stimulated with 180°-opposed sites of opto-PI3K activation show stable symmetry breaking, but cells stimulated with two 90°-opposed, frontward-oriented sites of opto-PI3K activation show frequent switching between the two directions. See **Movie S4** for an animated version of this data. Scale bars: 50 µm.

**Fig. 3.**
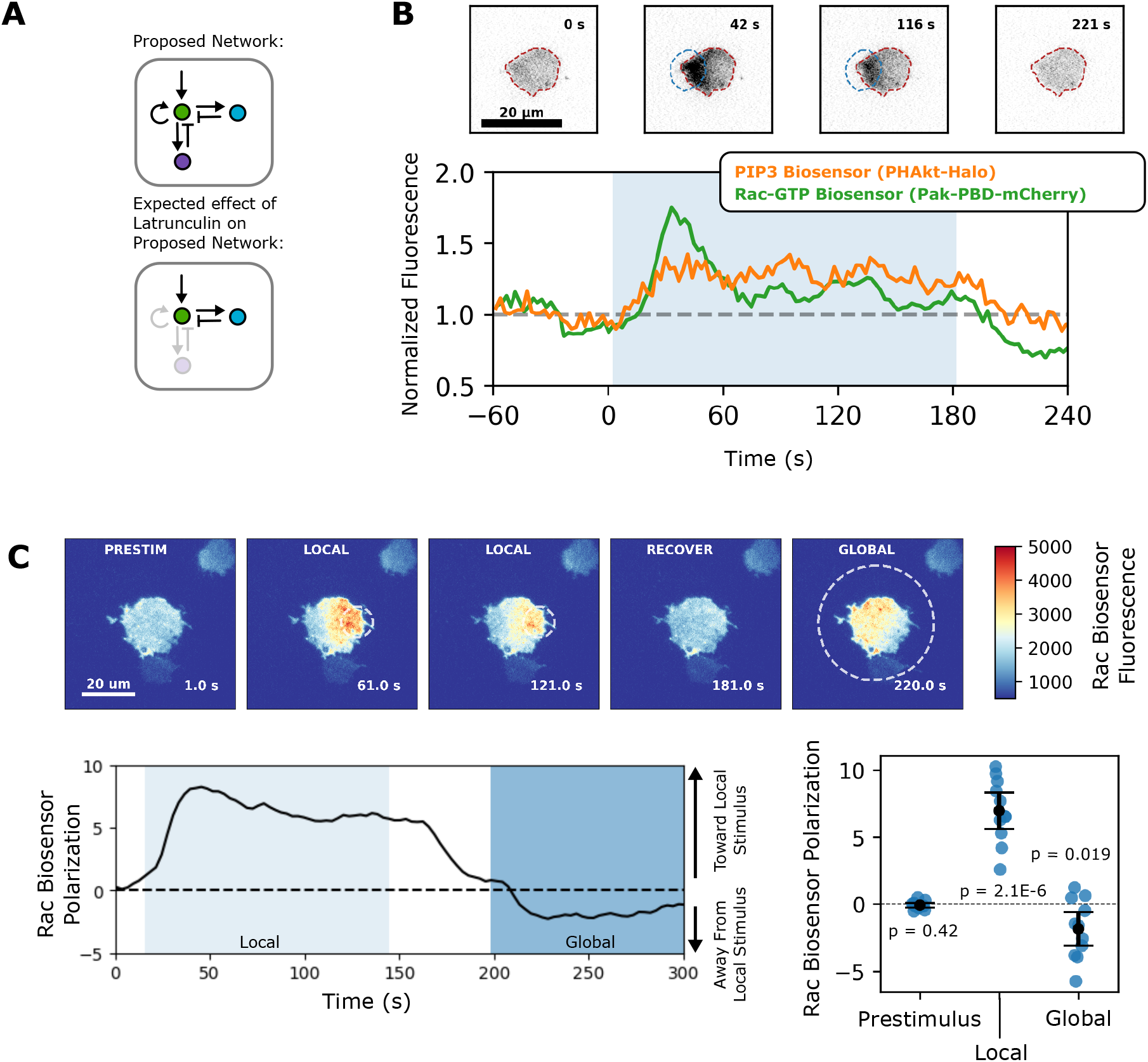
Shadow experiment provides direct evidence of local inhibition of Rac activation **A)** Inhibiting actin assembly with latrunculin treatment is expected to interrupt both local positive and global negative feedback. This disrupts front signal organization, but this organization can be restored with optogenetic activation of a leading edge signal like PI3K. This approach can be used to test for the presence of a PIP_3_-activated local inhibitor of Rac activation. **B)** Local recruitment of PI3K on one side of a latrunculin-treated cell shows local and transient Rac activation. This suggests that an actin-independent inhibition process is present in this context. **C)** To test the spatial range of this inhibition, we first stimulated one side of the cell to elicit a local transient Rac response, then briefly removed the input before globally activating PI3K. This revealed a shadow in which the pre-stimulated portion of the cell showed a weaker response than the naive portion of the cell, demonstrating local Rac inhibition at the site of first stimulation. See **Movie S5** for an animated version of this data.

We estimated the average angular velocity (in rotations per minute) of cells during each of the three phases of the assay across replicates of this assay **(Fig. 5C; Movie S7)**. We found statistically significant changes in these velocities **(Fig. 5D)**. The p values shown indicate the likelihood of observing these distributions under the null hypothesis of an average angular velocity of zero. We verified that the PIP_3_ biosensor matched the expected temporal profile in Fig. 5A on the population level **(Fig. 5E)**. Though the left and right sides of migrating cells reach the same steady state, the rates of change differ between the two sides. Turning behaviors of the cells coincide with the temporal changes in the input signal rather than the absolute levels of the signal. As further validation, we measured initial rates of PIP_3_ accumulation (as assayed via PHAkt recruitment) on each side for individual cells during the assay. In most cases, the initial rates matched the expected behavior **(Fig. S5)**.

By performing an experiment where we subject cells to local temporal changes in PIP_3_ in the absence of absolute differences in PIP_3_ across the cell, we demonstrate the sensitivity of cells to local signaling changes for cell guidance, consistent with a pilot pseudopod model of gradient interpretation.

## Discussion

To navigate toward sites of injury and inflammation, neutrophils must balance the decisiveness required to move in a single direction with the flexibility required to dynamically update this direction when appropriate. Previous work has shown that cells use local positive feedback (7–9, 55) and global negative feedback (5, 10, 11, 42, 43) to consolidate protrusive activity in a single direction, but these mechanisms alone do not account for the flexibility of directional orientation (3). Our current work suggests that a Rac-dependent local inhibitor plays a key role in neutrophil guidance by enabling this flexibility.

We leverage our ability to direct cell migration with optogenetically controlled PI3K to compare the relative responsiveness of different regions of polarized cells and to ask which spatiotemporal features of PIP_3_ signals inform cell orientation in a two-dimensional environment. Inhibiting both the global negative and local positive feedback loops that organize cell polarity, we use our optogenetic approach to provide direct evidence of a local inhibitor of Rac activation (as seen in our “shadow” experiments, Fig. 3). The activity of this inhibitor could explain how migrating cells avoid the polarity-locking behavior expected from local positive feedback and global inhibition alone (3). By controlling the dynamics of PIP_3_ accumulation in this same context, we demonstrate that a Rac-dependent negative feedback loop enables cells to detect the local rate of change of PIP_3_ **(Fig. 4)**. Taking into account reduced local responsiveness at the backs of cells **(Fig. 2)** (consistent with earlier studies (21, 23, 26–29)), we leverage our optogenetic inputs and an imaging-based control system to present cells with inputs that generate local changes in PIP_3_ in the absence of spatial differences in PIP_3_ **(Fig. 5)**. These experiments demonstrate that cells can decode the local rate of stimulus change for cell guidance.

**Fig. 4.**
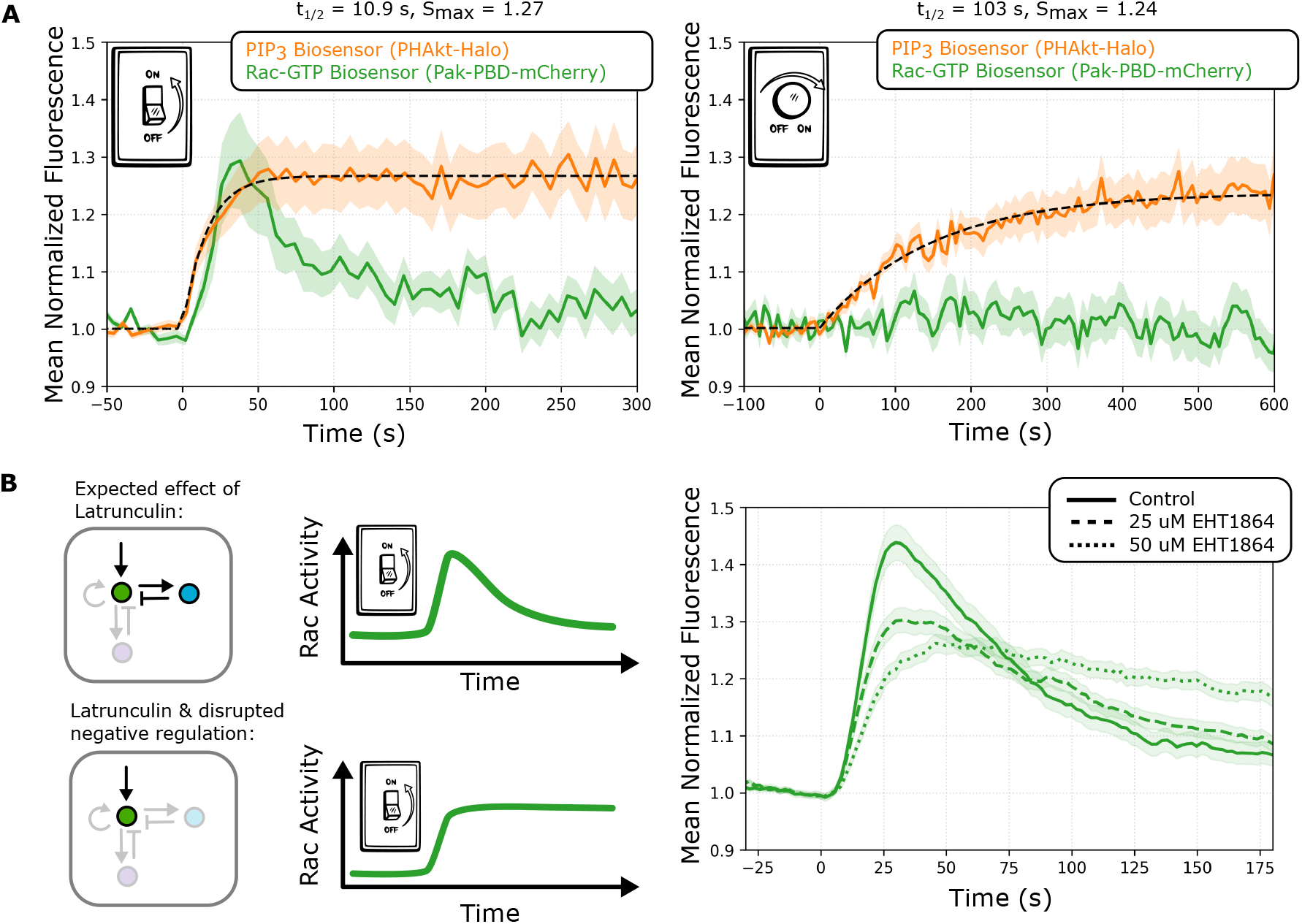
Local inhibition sensitizes cells to temporal changes in input signals via Rac-dependent negative feedback **A)** Exposing optoPI3K cells to a sudden increase in blue light causes transient Rac activation despite sustained PIP_3_ generation. In contrast, a slowly increasing ramp of PIP_3_ fails to elicit a strong Rac response even when similar PIP_3_ levels are generated to the step input protocol (left). The difference in responses to the rate-of-change of the input is consistent with adaptation via a negative feedback loop. (44, 45) S_max_ and t_1/2_ refer to the saturation and half-times to saturation respectively for the fit PIP_3_ curves. **B)** Pharmacologically inhibiting the ability of Rac to signal to downstream signaling partners shifts the dynamics of the Rac response toward a less adaptive and more linear response regime. These data suggest a Rac-dependent negative feedback loop.

**Fig. 5.**
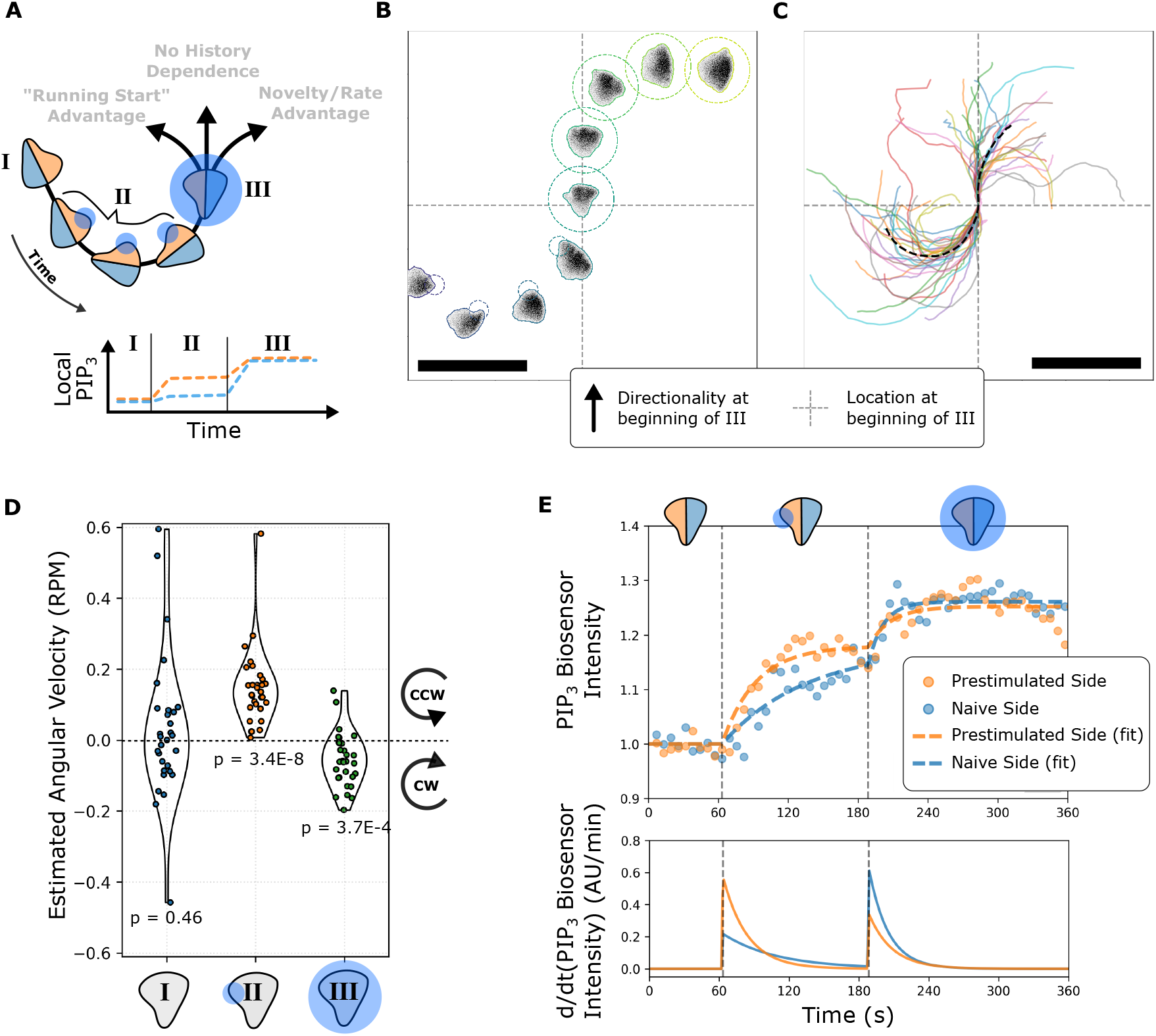
Cell fronts respond to local temporal changes in input signals **A)** To probe how cells decode the spatial and temporal features of input signals, we designed a test of the pilot pseudopod model by providing cells with different rates of change on their left and right sides despite the same absolute amount of signal on both sides. Cells are first tracked in I, then continuously stimulated (via opto-PI3K) at their left edge in II, and finally exposed to global illumination at III. This is expected to cause both sides of the cell to arrive at the same level of PIP_3_ signals with different histories. **B)** A tracked and stimulated cell responding to the stimulus scheme described in A. See **Movie S6** for an animated version of this subpanel. Scale bars: 50 µm. **C)** Tracks of many cells responding to the stimulus scheme described in A show a net bias towards reversal following global stimulation. See **Movie S7** for an animated demonstration of this behavior. Scale bars: 50 µm. **D)** Angular velocities were estimated for cells at the beginning of each of the phases described in A. p values shown describe a t-test for whether the mean of each distribution is zero. Cells transition from counterclockwise motion to clockwise motion following the switch from local stimulation (phase II) to global stimulation (phase III). **E)** The PIP_3_ signal on the left and right sides of migrating cells was calculated and fit with a time-delayed saturable function. As expected, both sides approach the same steady state at the end of phase III, but their histories differ. The changes in cell directionality correlate with the local derivative of input signals rather than the absolute levels of the input signal, consistent with the pilot pseudopod model of gradient interpretation.

**Fig. 6.**
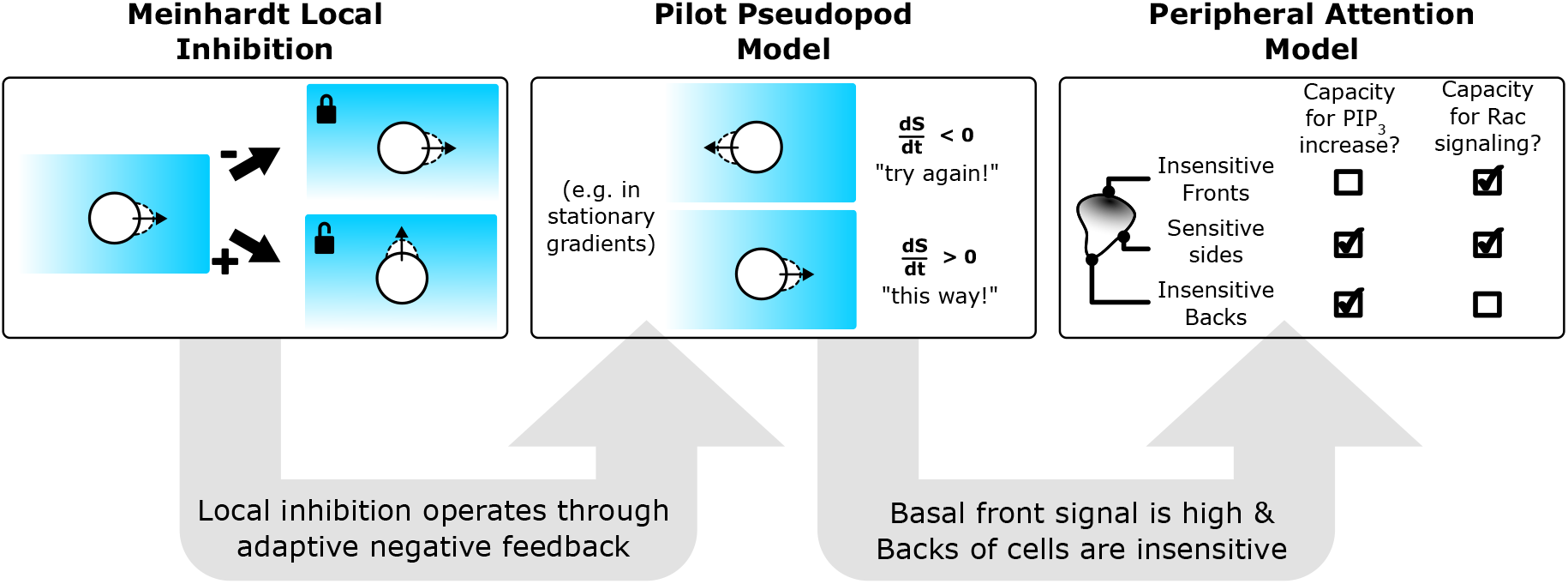
Cells interpret gradients using a modified pilot pseudopod program In this work, we have found direct evidence for Meinhardt-style local inhibition operating on the PI3K to Rac signaling axis. Furthermore, we have presented evidence that this local Rac negative feedback loop links the rate of change of input signals like PI3K signaling to the magnitude of Rac activation. This rate sensitivity is an expectation of the pilot pseudopod model. In the more complex case of a migrating cell, we find that the principles of the pilot pseudopod model apply with two modifications: the first being that the backs of cells are relatively insensitive to stimulation (as previously observed Arrieumerlou and Meyer 2005; Yoo et al. 2010; Nakajima et al. 2014; Hadjitheodorou et al. 2021); the second being that there exists a basal front-to-back gradient of PIP_3_ in migrating cells. Taken together, these two modifications make the lateral edges of migrating cells highly sensitive to increases in PIP_3_, implicating local inhibition in the ability of cells to turn and refine their direction during chemotaxis.

Our observations are consistent with the classic pilot pseudopod model, in which the rate of change of stimulus (rather than the absolute concentration of stimulus) is used to influence pseudopod lifetimes for cell guidance during chemotaxis (25). We extend this model to account for our observations of polarized signal sensitivity (similar to (26)). Since Rac responds to changes in PIP_3_, high basal signal at the fronts of cells renders them less sensitive than adjacent regions that have a larger capacity for increases in PIP_3_. The local negative feedback at the fronts of cells keeps the front from locking and enables cells to turn toward lateral edges experiencing temporal increases in signal. In much the same way that signals in peripheral vision direct the center of visual attention (56–58), temporal changes in signal at the periphery of the cell front alter its directionality.

The sensitivity to temporal input cues however does not extend to the backs of cells **(Fig. 2)** (21, 23, 26–29). Functionally, this insensitivity may help cells to integrate directional choices from the front over time through contractility and actin flow-dependent mechanisms (59, 60). This integration may be particularly important in shallow or changing gradients that may be too “noisy” for cells to accurately interpret instantaneously. For example, in shallow gradients, *Dictyostelium* (61) and zebrafish neutrophils (60) tend to orient themselves through gradient-biased left and right adjustments rather than through more dramatic reversals. In these contexts and in our work, cells maintain a rough heading while also retaining the ability to fine-tune directionality through these left-right “peripheral” adjustments.

The hybrid spatiotemporal guidance strategy we have described stands in contrast to systems of gradient interpretation that rely on only temporal cues, like those of bacteria. Because of their small size and rapid movement, bacteria use a purely temporal mechanism to migrate towards chemoattractants (48). Eukaryotic cells are larger and slower and take advantage of spatial signal processing to interpret guidance cues (62). However, eukaryotic cells do not solely decode gradients spatially but appear to also incorporate temporal information in their guidance (25). Importantly, the temporal information could either come from cells moving up fixed spatial gradients, or could come from dynamically evolving or even self-generated gradients (63–65). Experiments directly manipulating the spatiotemporal features of soluble chemoattractant gradients have shown that neutrophils reverse direction when they experience a temporal decrease in chemoattractant (66) and that cells require temporal increases in overall signaling during chemotaxis for efficient guidance (67). Our results suggest that these behaviors could be explained by the presence of a local Rac inhibitor at the fronts of cells and that studies examining cell behaviors in stable, dynamic, or self-generated gradients should consider both spatial and temporal features of these gradients as cells navigate them.

## Materials and Methods

### Cell culture

HL-60 cells were grown in RPMI 1640 media supplemented with l-glutamine and 25 mM Hepes (Mediatech) and containing 10% (vol/vol) heat-inactivated fetal bovine serum (Gibco). Cultures were maintained at a density of 0.2–1.0 million cells/ml at 37°C/5% CO2. HL-60 cells were differentiated by adding 1.5% (vol/vol) DMSO (Sigma-Aldrich) to actively growing cells and incubated for 5 days. HEK293T cells (used to generate lentivirus for transduction of HL-60 cells) were grown in DMEM (Mediatech) containing 10% (vol/vol) heat-inactivated fetal bovine serum and maintained at 37°C/5% CO2.

### Plasmids

Plasmids were assembled using a Golden-Gate-based modular cloning toolkit (68, 69).

The two key constructs for opto-PI3K system are based on the iLID system (31) and include a mTagBFP2-tagged, membrane localized, light sensitive component: AsLov2-SsrA-mTagBFP2-CAAX and an EGFP-tagged translocatable PI3K-binding component: ISH2-EGFP-SspBMicro. Upon 470 nm light exposure, the translocatable component localizes to the plasma membrane and the ISH2 domain recruits endogenous PI3K to the plasma membrane (8, 37).

Biosensors for PIP_3_ (32–34) and active Rac (35) were also created using the modular cloning kit and were designed to be continuously imaged during experiments without activating the blue-light-sensitive optogenetic system. For experiments shown in this paper, PIP_3_ was detected using a Halotagged (70) PH domain from Akt labeled with Janelia Fluor 646 (71), and active Rac was detected using an mCherry-tagged Pak-PBD domain from Pak1.

### Transduction of HL-60 cells

HEK293T cells were seeded into six-well plates and grown until about 80% confluent. For each well, 1.5 µg pHR vector (containing the appropriate transgene), 0.167 µg vesicular stomatitis virus–G vector, and 1.2 µg cytomegalovirus 8.91 vector were mixed and prepared for transfection using TransIT-293 transfection reagent (Mirus Bio) per the manufacturer’s instructions. After transfection, cells were grown for an additional 3 d, after which virus-containing supernatants were harvested and concentrated 20-fold using a Lenti-X Concentrator (Takara Bio Inc.) per the manufacturer’s instructions. Concentrated viruses were frozen and stored at −80°C until needed. For all transductions, the thawed virus was mixed with about 0.3 million cells in growth media supplemented with polybrene (8 µg/ml) and incubated overnight. Cells expressing desired transgenes were isolated by FACS.

### Preparation of cells for microscopy

Cells expressing Halo fusion proteins were stained with 10 nM JF646 (Janelia) for 10-15 minutes at 37°C in complete media and then rinsed once with complete media (RPMI with 10% FBS) before placing cells in reduced-serum media for migration assays. We used an under-agarose preparation (72) with reduced serum (RPMI with 2% serum) to keep cells adjacent to the coverglass for TIRF imaging and optogenetic stimulation. This preparation involved layering 2-2.5% low melting point agarose onto cells after they had been allowed to attach to the glass for 10-15 minutes.

### Microscopy Hardware

Hardware used for the experiments included an Eclipse Ti inverted microscope equipped with a motorized laser TIRF illumination unit, a Borealis beam-conditioning unit (Andor Technology), a 60× Plan Apochromat TIRF 1.49 NA objective (Nikon), an iXon Ultra electron-multiplying charge-coupled device camera, and a laser merge module (LMM5; Spectral Applied Research) equipped with 405-, 488-, 561-, and 640-nm laser lines. All hardware was controlled using Micro-Manager (36, 38) (University of California, San Francisco), and all experiments were performed at 37°C and 5%CO2.

Activity of opto-PI3K was controlled via a 470-nm (blue) LED (Lightspeed Technologies) that transmitted light through a custom DMD (Andor Technology) at varying intensities by connecting the LEDs to the analog outputs of a digital-to-analogue converter and setting the LED voltages using serial commands via custom Python code. Our microscope is equipped with two stacked dichroic turrets such that samples can be simultaneously illuminated with LEDs using a 488-nm longpass dichroic filter (Chroma Technology Corp.) in the upper turret while also placing the appropriate dichroic (Chroma Technology Corp.) in the lower turret for TIRF microscopy.

### Automated microscopy and light-activation pipeline

Images were collected, and light was spatially patterned using customized tracking and light-patterning code written in python and making use of Pycromanager (40). The code we used for our experiments and analysis is available on GitHub (https://github.com/weinerlab).

In brief, a program scanned locations in a prepared well while running a segmentation algorithm at each location. If a segmented object was found to be an appropriate size and intensity, the program would then image that object for 30 seconds to test whether it was moving at a speed appropriate for a migrating cell. If so, it moved that object into the center of the field of view and initiated a tracking and stimulus protocol to track the cell over time and deliver programmed stimuli described in this paper.

### Image and Data Analysis

#### Segmentation and background correction

Collected TIRF images were segmented based on fluorescence intensity from the PHAkt channel. Background corrections were accomplished for each channel by first segmenting the backgrounds of images, then fitting either a plane

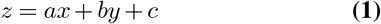

(for 400px-400px cropped images) or a two-dimensional quadratic equation

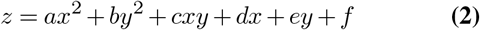

(for full size 1024px-1024px images) to the selected background pixel intensities. After background subtraction, if a cell’s average intensity was less than 200 units, it was excluded from analysis due to unreliable baseline normalization.

#### Global biosensor quantification

For quantification of biosensor dynamics in single cells, we first segmented single cells and background-subtracted the images. In some cases, cells were too dense to segment accurately, so we instead estimated the centers of cells by blurring the image and using a peak finding algorithm. We then used a 20 by 20 pixel (4.3 by 4.3 µm) square centered at those locations as a sampled approximation of whole-cell dynamics. We next took background-subtracted single cell average traces and divided each one by the baseline pre-stimulus intensity (we excluded cells with a baseline background-subtracted fluorescence below 100 units to avoid errors from dividing by small numbers). These normalized traces were then averaged together. Corresponding timestamps were extracted from Micromanager data.

#### Classification of movement toward frontward stimuli in two-spot assays

The distance traveled toward each potential site was calculated and classified as going toward either the frontward stimulus, the rearward stimulus, or neither if the cell did not move more than 20 µm in either direction. These “neither” cells were frequently nonresponders that appeared to have lost expression of one or both of the optogenetic components.

#### Fold-change in biosensor recruitment as function of distance from front

The front of the cell was defined as the point on the boundary of the segmented region that intersected with the vector describing the direction of motion of the cell. After background correction, all pixels were binned based on their distance from this point and averaged. Shown in Fig. 2F is data from 39 cells. The initial front-back pre-stimulus distributions were calculated and compared to the post-stimulus distribution to generate this data.

#### Detection of reversals in two-spot assays

To detect reversals in directionality during the two-spot assays shown in **Fig. 2G**, we first slightly smoothed the cell trajectories in x and y using a Savitzky-Golay filter (window length of 9 and polynomial order 3) and then calculated the angle and displacement between successive frames. We disregarded angles from timepoints where the displacement was less than 2 µm/min as these were unreliable measurements. Instead, we assume these values were unchanged from the last confidently measured angle. Finally, we “unwrapped” the values, which accounts for the periodicity of angular measurements, converting the values into accumulative measurements by adding or subtracting multiples of 2π where necessary. For each cell, we then classified movement as being aligned with a stimulus if it was within 0.4 radians of the angle of that stimulus (in either direction). If cells spent any time classified as being aligned with one stimulus and the other during the same continuous period of stimulation, they were considered reversers.

#### Rac biosensor polarization metric

As a metric for calculating the directionality of the Rac biosensor distribution in Fig. 3, we found the vector given by the difference between the unweighted center of mass of the segmented cell and the center of mass weighted by the fluorescence signal in each frame during the local to global stimulation protocol. We then took the magnitude of the component of this vector in the direction of optogenetic stimulation and subtracted the average pre-stimulus values to account for differences in illumination or coverslip proximity. Values shown in the graph are approximately when directional signals peaked on average (50 s post-stimulus).

#### Fitting curves to PIP_3_ accumulation dynamics

To estimate half-times for PIP_3_ accumulation, we fit a piecewise dynamic model to the population-averaged, background-subtracted, and baseline-normalized fluorescence data using scipy’s curve fit function. The equation was

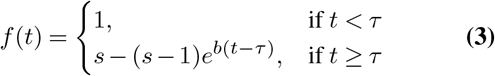

where s is the saturation value, tau represents an adjustable delay, and b is related to the half-time:

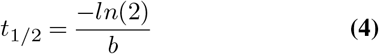

Similarly, for the curves shown in **Fig. 5**, we used a two-stage saturable function:

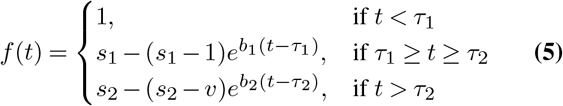

where

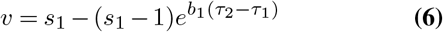

#### Estimation of angular velocity

To estimate angular velocity of continuously-turning cells shown in Fig. 5D, we processed the angular movement of cells as described above for detection of reversals in two-spot assays. We then fit a line to the angle values as functions of time, recorded the slope values, and converted them into revolutions per minute. For pre-stimulus velocities (I), the entire one-minute pre-stimulus time period was used. For the local stimulus velocities (II), the two minute period immediately preceding the switch to global stimulation was used. For the global stimulus velocities (III), the two minute period immediately following the switch to global stimulation was used.

#### Spatial dynamics of PHAkt dynamics and estimation of initial rates

To estimate the PIP_3_ dynamics on the left and right sides of cells as they were stimulated according to the schematic in **Fig. 5A**, we calculated the cell direction and used that as an axis to define the left and right sides of the cells. We calculated the background-corrected mean fluorescence signals on each side over time and averaged the globalbaseline-corrected dynamics to produce Fig. 5E. We fit an equation to these dynamics to better estimate the rates of accumulation on each side. We estimated rates for individual cells on their left and right halves by fitting a line to the signal during the first 30 s of stimulus response.

### Measurement of Rac-GTP by GLISA

For biochemical verification of Rac-GTP dynamics in response to acute optogenetic activation of PIP_3_, we used a colorimetric Rac 1,2,3-GLISA (Cytoskeleton Inc).

After testing several cell densities to determine the linear range of the assay, we settled on using 2E5 undifferentiated HL-60 cells per sample. Following this, 2E5 cells in 10 uL DPBS were placed in centrifuge tubes and exposed to a saturating dose of blue LED light for varying amounts of time. Cells were lysed in 120 µL ice cold GL36 buffer and 100 µL lysate was transferred into ice-cold tubes and immediately snap-frozen in liquid nitrogen. The remaining 20 µL were used to verify that each of the samples had similar protein concentrations (of approximately 0.16 mg/mL).

Once all samples were collected, the 100 uL aliquots were thawed in a room temperature water bath and processed according to the manufacturer’s instructions.

## ACKNOWLEDGEMENTS

We thank Anne Pipathsouk and Brian Graziano for helpful discussion, Kirstin Meyer, Ben Winer, and Tamas Nagy for a critical reading of the manuscript, and all members of the Weiner lab for their support. The JF646 HaloTag dye was kindly provided by Dr. Luke Lavis. This work was supported by an NSF Predoctoral Fellowship (JT), National Institutes of Health grant GM118167 (ODW), National Science Foundation/Biotechnology and Biological Sciences Research Council grant 2019598 (ODW), the National Science Foundation Center for Cellular Construction (DBI-1548297, ODW), and a Novo Nordisk Foundation grant for the Center for Geometrically Engineered Cellular Systems (NNF17OC0028176, ODW).

## Supplementary Information

**Movie S1:**

PIP_3_ and Rac biosensors are locally recruited in response to local blue light stimulation.

**Movie S2:**

Migrating cells respond to single-spot assays by altering their directionality. The live-cell fluorescent Rac biosensor (Pak-PBD-mCherry) is shown. Scale bars are 50 µm.

**Movie S3:**

Computer-controlled local activation can continuously turn migrating cells. The live-cell fluorescent Rac biosensor (Pak-PBD-mCherry) is shown. X and Y axes units are µm.

**Movie S4:**

Migrating cells respond to two-spot assays differently depending on the positioning of the spots. 180°-opposed spots create a winner-take-all scenario, while 90°-opposed, frontward-oriented spots cause cells to switch between the two directions. The live-cell fluorescent Rac biosensor (Pak-PBD-mCherry) is shown. Scale bars are 50 µm.

**Movie S5:**

A local-to-global stimulation assay shows direct evidence of local inhibition of Rac. The live-cell fluorescent Rac biosensor (Pak-PBD-mCherry) is shown in latrunculin-treated cells (10 µM).

**Movie S6:**

Migrating cell responding to local temporal changes in input signal. The live-cell fluorescent Rac biosensor (Pak-PBD-mCherry) is shown. Scale bar is 50 µm.

**Movie S7:**

Multiple migrating cells responding to local temporal changes in input signal. The live-cell fluorescent Rac biosensor (Pak-PBD-mCherry) is shown. X and Y axes units are µm.

**Fig. S1:**
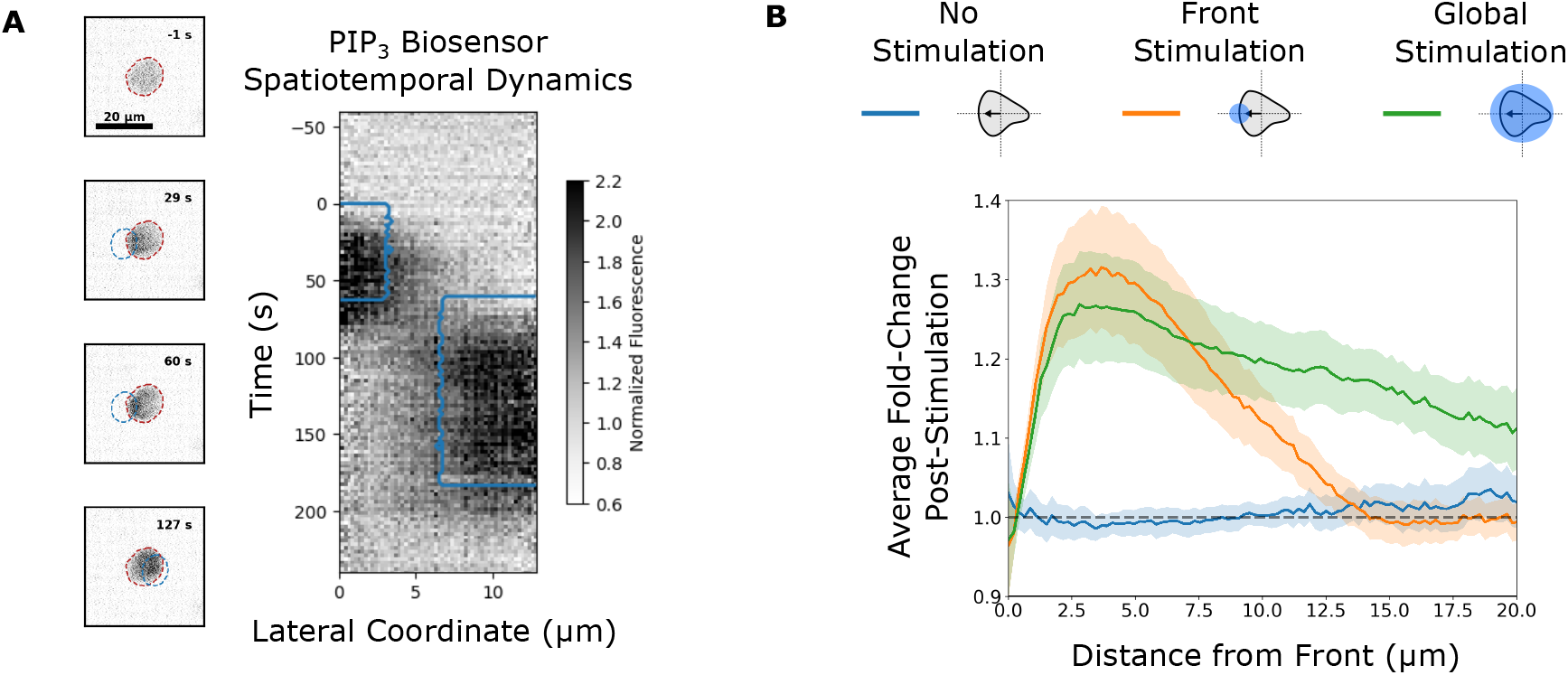
Spatial control of PIP_3_ production using opto-PI3K. **A)** TIRF microscopy of a latrunculin-treated (10 µM) cell that was exposed to blue light, first on its left side and then on its right. Shown on the right is a depiction of the PIP_3_ biosensor fluorescence signal (PHAkt-Halo (JF646)) as a function of time and the lateral position of pixels. The fluorescence signal closely corresponds in time and space to the blue-light activation regions, shown with the blue outline. See textbfMovie S1 for an animated version. **B)** The average fold-change in PIP_3_ biosensor distributions along the front-back axis in migrating differentiated cells following blue light stimulation. Front-localized blue light stimulation (orange line) elicits local increases in the biosensor specifically at the fronts of cells.

**Fig. S2:**
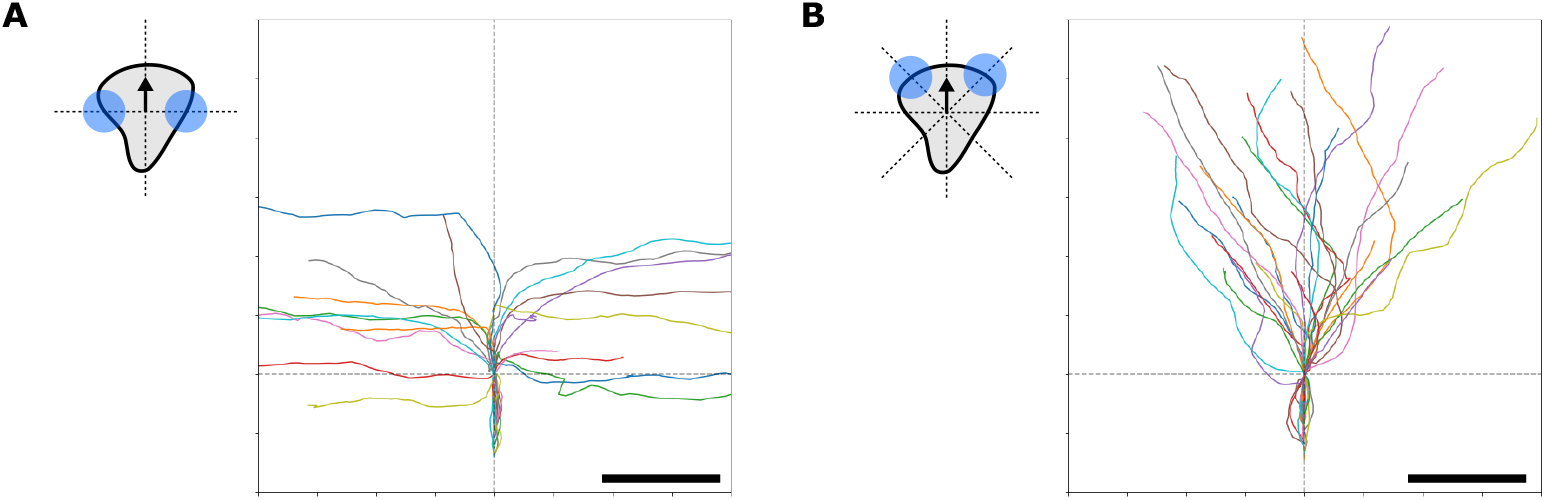
Cell paths in two-spot assays **A)** Cells stimulated simultaneously at opposite lateral edges stably move in only one of the two directions separated by 180°. **B)** Cells stimulated simultaneously at two sites near the front (separated by 90°), sometimes show switching behaviors, first following one site of opto-PI3K activation and then the other.

**Fig. S3:**
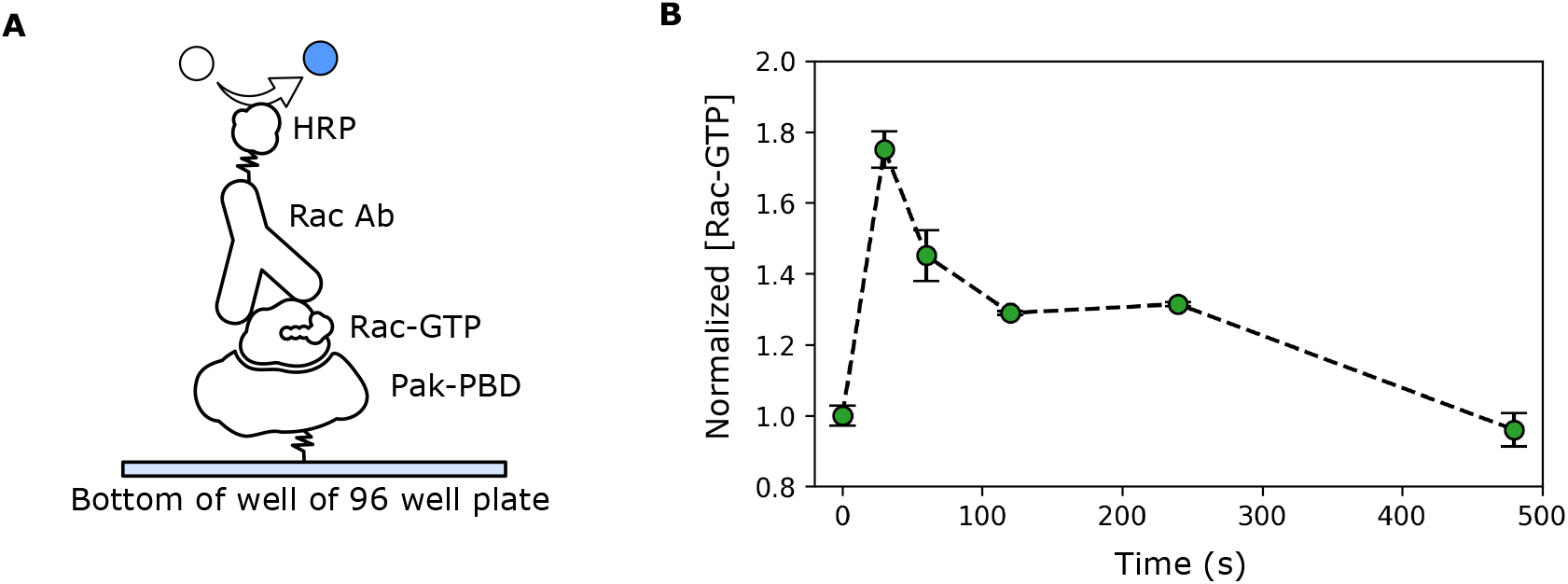
Confirmation of live-cell Rac biosensor dynamics with a Rac GLISA. **A)** Schematic of the ELISA-like mechanism used in the assay. Plate-bound Pak-PBD binds only the active form of Rac (Rac-GTP). This can then be detected and quantified through a colorimetric assay using an HRP-conjugated Rac antibody. **B)** GLISA-based measurements of relative Rac-GTP in blue-light-exposed opto-PI3K cells closely matches the dynamics observed using the live-cell Rac biosensor, Pak-PBD-mCherry (compare with **Fig. 4A, Left**)

**Fig. S4:**
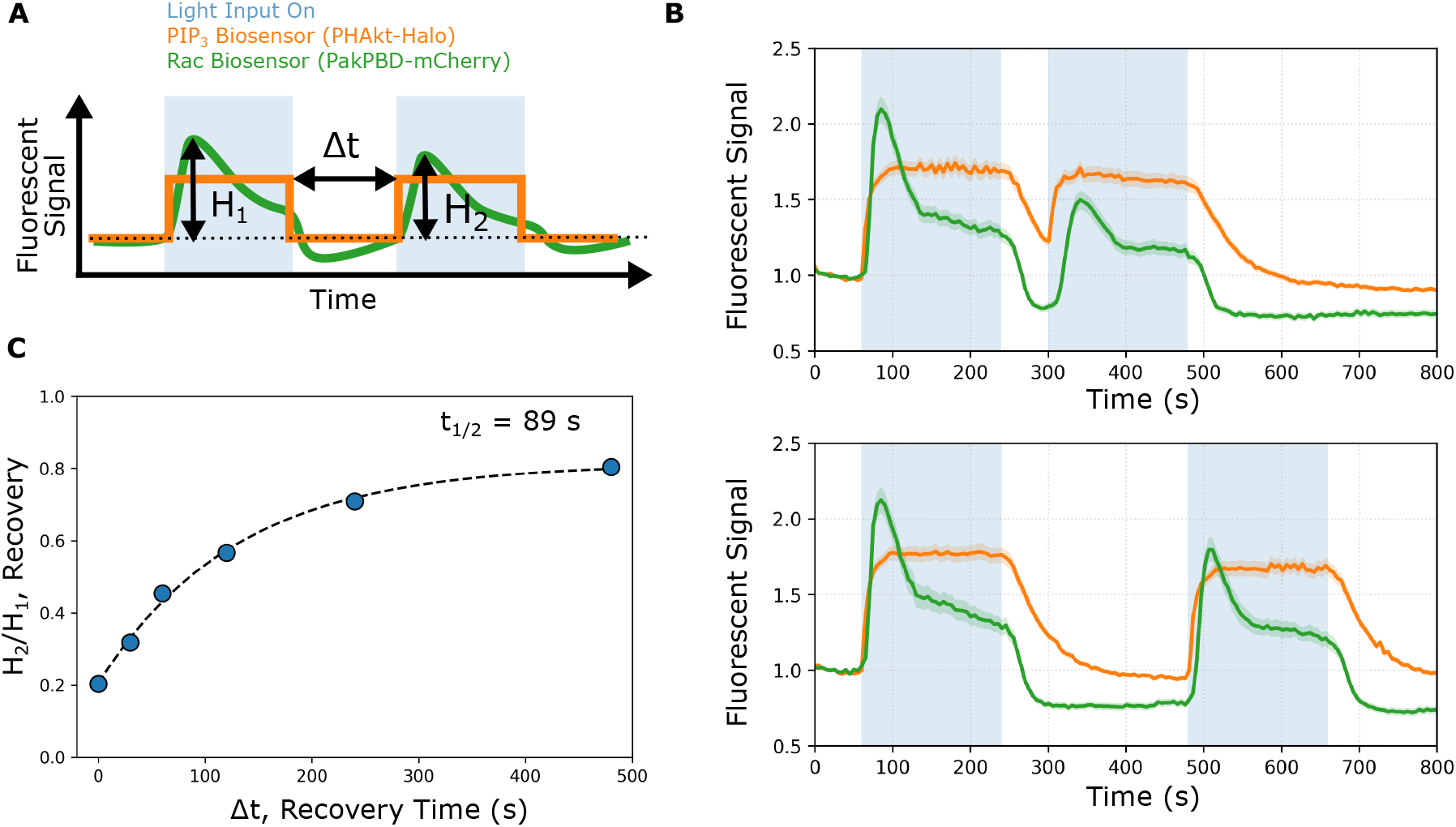
Timescale of reversibility of Rac inhibition. **A)** Experimental schematic: Latrunculin-treated (10 µM) cells were exposed to two pulses of blue light separated by a variable amount of recovery time. We then calculated the ratio of the heights of the peaks of the first and second Rac responses. **B)** Two example curves showing the response of the PIP_3_ and Rac biosensors to two pulses of blue-light activation with recovery times of 60 seconds (top) and 240 seconds (bottom). **C)** Degree of recovery as a function of recovery time. The response has a recovery half-time of 89 seconds.

**Fig. S5:**
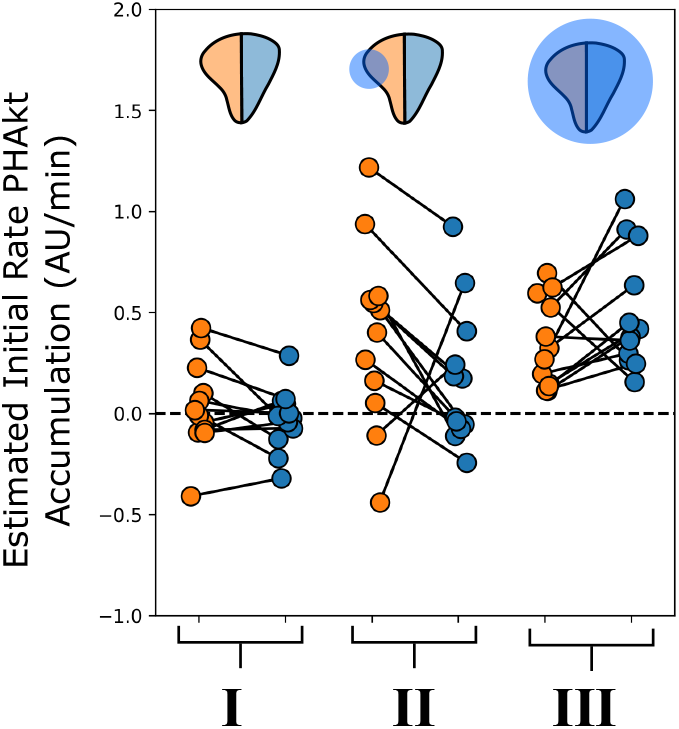
Local PIP_3_ dynamics in single migrating cells during reversal assay: Differences in the estimated initial rates of PHAkt-Halo accumulation on the left and right sides of cells during the reversal assay **(Fig. 5)**. As expected, the majority of cells show similar dynamics on their left and right sides pre-stimulation (phase I). However, cells show higher rates of increase in PIP_3_ on the left side during the local stimulation phase of the assay (phase II) and on their right sides during the subsequent global stimulation phase of the assay (phase III).

